# DHX15 regulates CMTR1-dependent gene expression and cell proliferation

**DOI:** 10.1101/113209

**Authors:** Francisco Inesta-Vaquera, Alison Galloway, Laurel Chandler, Alejandro Rojas-Fernandez, Viduth K. Chaugule, Simone Weidlich, Mark Peggie, Victoria H Cowling

## Abstract

CMTR1 contributes to mRNA cap formation by methylating the O-2 position of the 1^st^ transcribed nucleotide ribose. mRNA cap O-2 methylation has roles in mRNA translation and self-RNA tolerance in innate immunity, however its role in cell physiology is unclear. We report that CMTR1 is recruited to Serine-5 phosphorylated RNA Pol II CTD, facilitating cotranscriptional methylation. We isolated CMTR1 in a complex with DHX15, an RNA helicase functioning in splicing and ribosome biogenesis, and characterised it as a regulator of CMTR1. When bound to DHX15, CMTR1 activity is repressed and prevented from binding to RNA pol II, thus constraining 1^st^ nucleotide methylation to a co-transcriptional event. Conversely CMTR1 activates DHX15 helicase activity and influences its nuclear localisation, which is likely to impact on several nuclear functions. The impact of the CMTR1-DHX15 interaction is complex and will depend on the relative expression of these enzymes and their interactors, and the cellular dependency on different RNA processing pathways. In HCC1806 cells, the DHX15-CMTR1 interaction controls ribosome loading of a subset of mRNAs and impacts on cell proliferation.

## Introduction

Formation of the mRNA cap initiates the maturation of RNA pol II transcripts into translation-competent mRNA (Furuichi, 2015). The mRNA cap protects transcripts from degradation and recruits protein complexes involved in nuclear export, splicing, 3’ processing and translation initiation (Ramanathan et al, 2016; Topisirovic et al, 2011). mRNA cap formation initiates with the addition of an inverted guanosine group, via a tri-phosphate bridge, to the first transcribed nucleotide of nascent RNA pol II transcripts. Subsequently, this guanosine cap is methylated on the N-7 position to create the cap 0 structure, which binds efficiently to CBC, eIF4F and other complexes involved in RNA processing and translation initiation. The initial transcribed nucleotides are further methylated at several other positions, in a species-specific manner. In mammals, the O-2 position of the riboses of the 1^st^ and 2^nd^ transcribed nucleotides are sites of abundant methylation (Langberg & Moss, 1981).

A series of enzymes catalyses mRNA cap formation, which have different configurations in different species (Shuman, 2002). In mammals, RNGTT/CE catalyses guanosine cap addition and RNMT-RAM catalyses guanosine cap N-7 methylation. RNGTT/CE and RNMT-RAM are recruited to RNA pol II at the initiation of transcription (Buratowski, 2009). CMTR1 and CMTR2 methylate the O-2 position of 1^st^ and 2^nd^ transcribed nucleotides riboses, respectively (Belanger et al, 2010; Werner et al, 2011).

CMTR1 (ISG95, FTSJD2, KIAA0082) was first identified as a human-interferon regulated gene. It was recognised to have several functional domains including a methyltransferase domain (Haline-Vaz et al, 2008). Subsequently, CMTR1 was biochemically characterized as the O-2 ribose methyltransferase of the first transcribed nucleotide (Belanger et al, 2010). Recently, the catalytic domain of CMTR1 was crystalized with s-adenosyl methionine and a capped oligonucleotide (Smietanski et al, 2014). Our knowledge of the biological function of CMTR1 and 1^st^ nucleotide O-2 methylation is scant. An early report in sea urchin eggs demonstrated that following fertilization N-7 and O-2 cap methylation is associated with translational upregulation of a subset of maternal transcripts (Caldwell & Emerson, 1985). During *Xenopus* oocyte maturation, 1^st^ nucleotide O-2 methylation significantly increases translation efficiency and is required for the translation of maternal mRNA (Kuge et al, 1998; Kuge & Richter, 1995). In mice, a significant proportion of 1^st^ nucleotides were found to be O-2 methylated on the ribose, although the relative proportion of this methylation varied between organs, indicating a regulated event (Kruse et al, 2011). O-2 methylation is likely to influence recruitment of cap binding complexes and promote ribosomal subunit binding (Muthukrishnan et al, 1976b).

Recently, the composition of 5' cap has been identified as an important determinant of self (host) versus non-self (viral) RNA during viral infection (Leung & Amarasinghe, 2016). The absence of O-2 methylation in viral transcripts was demonstrated to results in enhanced sensitivity to the interferon-induced IFIT proteins, which lead to the discovery that this first nucleotide methylation mark distinguishes self from non-self RNA (Daffis et al, 2010). Schuberth-Wagner *et al.,* provided the first evidence that CMTR1 is involved in the recognition of self RNA (Schuberth-Wagner et al, 2015). CMTR1-dependent O-2 methylation of the first nucleotide abrogated activation of RIG-I, a helicase that initiates immune responses on interaction with uncapped or aberrantly capped transcripts. CMTR1 depletion was able to trigger a RIG-I-dependent immune response similar to that triggered by viral transcripts (Schuberth-Wagner et al, 2015).

Here we report the first regulator of CMTR1 function. We report that CMTR1 and the DEAH-box RNA Helicase, DHX15, form a stable complex in cells and reciprocally influence each others activity and action. DXH15 restrains CMTR1 activity and localisation constraining first nucleotide methylation to a co-transcriptional event. CMTR1 impacts on DHX15 action by activating helicase activity and influencing nuclear localisation. Disruption of the CMTR1-DHX15 interaction leads to increased ribosome loading of a subset of mRNAs involved in key metabolic functions, which impacts on cell proliferation.

## Results

### CMTR1 interacts directly with DHX15

To investigate the regulation and function of CMTR1, we identified CMTR1-interacting proteins. HA-CMTR1 was immunoprecipitated (IP) from HeLa cell extracts using anti-HA antibodies (Fig 1a). IPs were resolved by SDS-PAGE and purified proteins identified by mass spectrometry. In HA-CMTR1 IPs, DHX15, a 95kDa DEAH box RNA helicase was identified (accession O43143), with significant mascot scores and protein coverage (Fig EV1a). In order to verify their interaction, GFP-CMTR1 and FLAG-DHX15 were transiently expressed in HeLa cells and co-immunoprecipitated from cell extracts (Fig 1b). In order to investigate the action of endogenous CMTR1, an antibody was raised against recombinant human CMTR1. In western blot analysis of HeLa cell extracts, the anti-CMTR1 antibody recognised an 110kDa band, the expected molecular weight of CMTR1, which was reduced following transfection of two independent CMTR1 siRNAs (Fig EV1b). IPs performed with the anti-CMTR1 antibody purified endogenous CMTR1 (Fig EV1c). The interaction of endogenous CMTR1 and DHX15 was confirmed by anti-CMTR1 and anti-DHX15 antibody IPs in HeLa, HCC1809, U2OS and HEK293 cell extracts, with species-matched IgG used as a negative control (Fig 1c). In *CMTR1* del cells, in which expression of guide RNAs and Cas9 ablated *CMTR1* expression, the anti-CMTR1 antibody was unable to purify DHX15, discounting non-specific interactions of DHX15 with resin or antibody (Fig1d). Of note, inhibition of CMTR1 expression did not impact on DHX15 expression and, conversely, inhibition of DHX15 expression did not impact CMTR1 expression (Fig1d and EV1b). Since CMTR1 and DHX15 bind RNA the dependency of their interaction on RNA was investigated (Bohnsack et al, 2009; Smietanski et al, 2014). The CMTR1-DHX15 interaction was sustained on RNaseA treatment and therefore is not maintained by an RNA connector (Fig 1e). RNaseA was confirmed to be functional by digestion of HeLa cell RNA (Fig EV1d). In HeLa cells, DHX15 and CMTR1 are expressed at equivalent molar ratio (Nagaraj et al, 2011). Equimolar recombinant human CMTR1 and His^6^-DHX15 co-immunoprecipitated, confirming their direct interaction (Fig 1f and EV1e).

**Figure 1.**
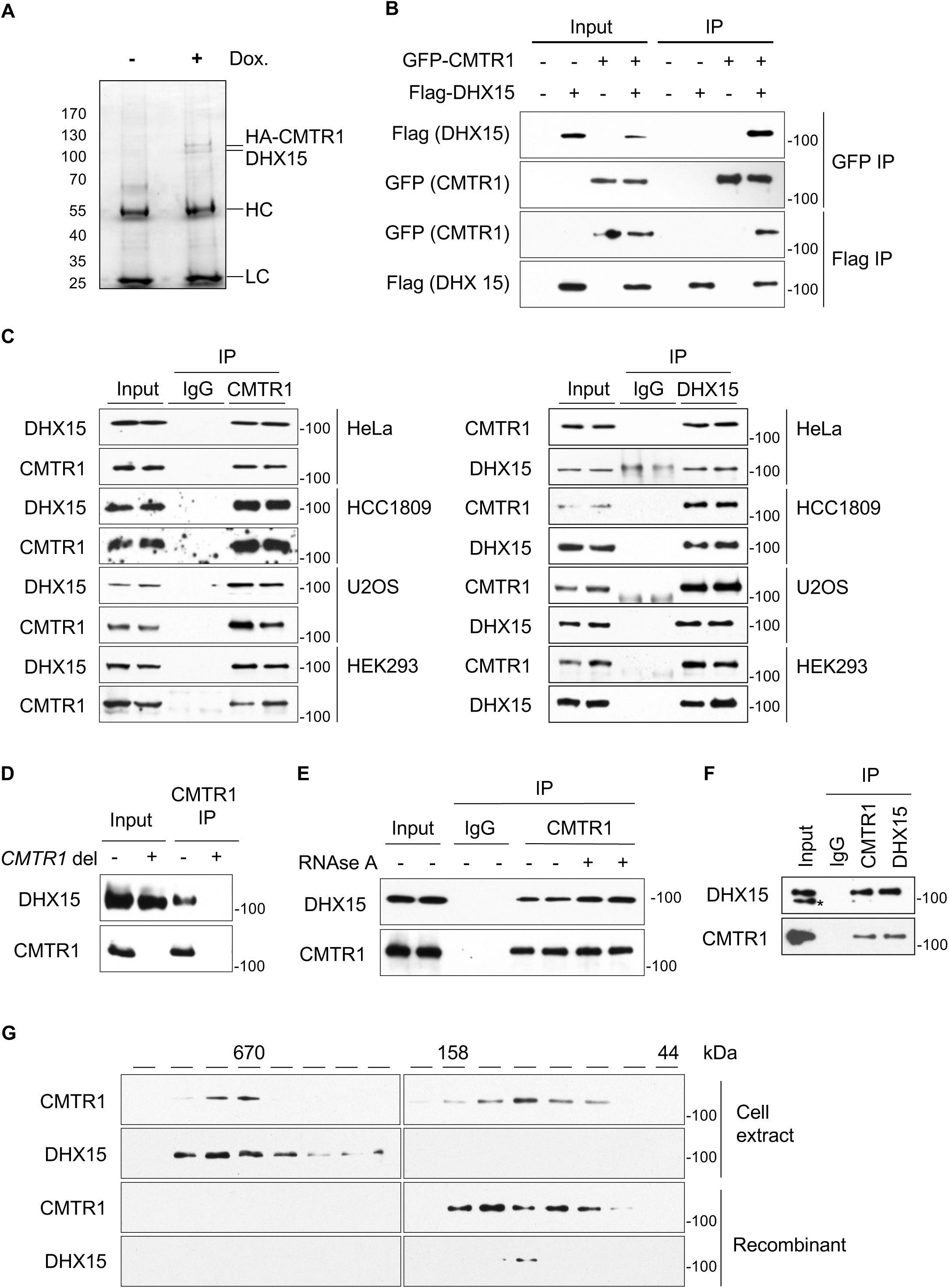
DHX15 interacts with CMTR1. a) Anti-HA antibodies immunoprecipitation (IP) of HA-CMTR1 following Doxycyline (Dox) induction in HeLa cells. IPs resolved by SDS-PAGE and Coomassie Blue-stained. HA-CMTR1, DHX15, antibody heavy chain (HC), light chain (LC) indicated. b) pcDNA5 GFP-CMTR1 and FLAG-DHX15 transiently expressed in HeLa cells. IPs performed on cell extracts using anti-GFP and FLAG antibodies prior to western blot analysis. c) Endogenous CMTR1 (left panels) or DHX15 (right panels) IPs from extracts of cell lines indicated prior to western blot analysis. Sheep IgG used for IP control. d) Endogenous CMTR1 IP performed on HeLa (-) or HeLa *CMTR1* del (+) cell extracts, prior to western blot analysis. e) CMTR1 IPs from HeLa cells untreated or incubated with RNAase A, prior to western blot analysis. f) 100ng recombinant CMTR1 and 100ng His^6^-DHX15 were mixed and subjected to IP with indicated antibodies, prior to western blot analysis. * denotes non-specific band. g) HeLa cell extract, recombinant CMTR1 or recombinant DHX15 resolved on a Superdex s200 10/30 column. Fractions analysed by western blot. Elution of standards indicated.

To gain insight into the relative abundance of CMTR1 and DHX15 complexes in HeLa cells, gel filtration analysis was performed (Fig 1g, upper panels). Recombinant CMTR1 and His^6^-DHX15 monomers were also analysed, which migrated as expected for ~100kDa monomers (Fig 1g, lower panels). In HeLa cell extracts, approximately half of CMTR1 and DHX15 eluted at >670kDa, which is likely to contain the CMTR1-DHX15 complex. Approximately half of cellular CMTR1 also eluted at 100kDa, consistent with monomeric CMTR1. Analysis of HA-CMTR1 IPs indicated that the predominant CMTR1-interacting protein is DHX15 (Fig 1a), and therefore in HeLa cells about half of cellular CMTR1 is in a DHX15-CMTR1 complex and half is in a CMTR1 monomer. Conversely, cellular DHX15 monomers were not observed, with DHX15 complexes peaking >670kDa and <670kDa, indicating two or more complexes. DHX15 is expected to migrate in several large complexes since it is also present in complexes associated with splicing and ribosomal biogenesis (Robert-Paganin et al, 2015)(Memet et al, 2017). The relative abundance of DHX15 and CMTR1 complexes is likely to be different in different cell lines, depending on their abundance and that of other co-factors.

### CMTR1 G-patch domain interacts with the DHX15 OB-fold

CMTR1 is an 835 residue protein possessing a nuclear localisation signal (residues 2-19), G-patch domain (residues 85-133), RrmJ-type SAM-dependent O-2 methyltransferase domain (residues 170-550), guanylyltransferase-like domain (residues 560-729), and WW domain (755-790) (Fig 2a) (Aravind & Koonin, 1999; Belanger et al, 2010; Haline-Vaz et al, 2008; Smietanski et al, 2014). To establish the regions of CMTR1 and DHX15 which interact, HeLa cells were transfected with N-terminal GFP-tagged CMTR1 wild type (WT) or mutants ∆N (25-835), ∆G (143-835), 1-143, Gp (85-143) or GFP alone (Fig 2a). Endogenous DHX15 co-immunoprecipitated with GFP-CMTR1 WT and mutants with the exception of GFP-∆G, the G-patch deletion mutant (Fig 2b). Furthermore, DHX15 co-immunoprecipitated with GFP-Gp, the G-patch domain of CMTR1 alone. GFP did not bind to DHX15.

**Figure 2.**
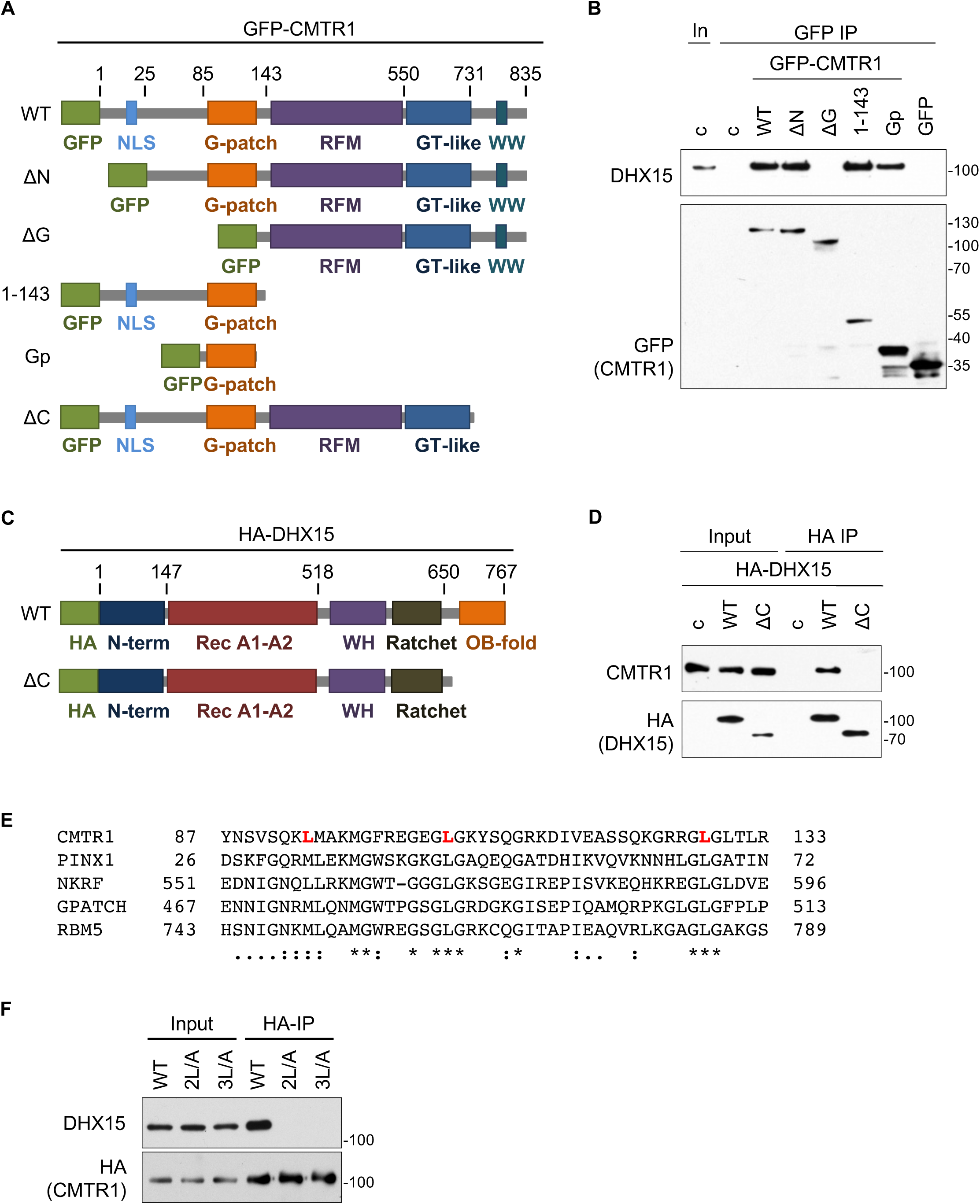
CMTR1 G-patch binds to DHX15 OB-fold. a) CMTR1 domains and mutants generated. b) HeLa cells transfected with pcDNA5 GFP, GFP-CMTR1 and mutants. Anti-GFP-antibody IPs analysed by western blot. In, cell extracts. c) DHX15 domains and mutants generated. d) HeLa cells transfected with pcDNA5, pcDNA5 HA-DHX15 and mutants. Anti-HA antibody IPs analysed by western blot. e) Alignment of G-patch domains in DHX15 interactors generated by Clustal Omega Alignment tool. Conserved leucines in CMTR1 in red. Residues:".", weakly similar; ":", strongly similar; "*", conserved. f) HeLa cells transfected with pcDNA5 HA-CMTR1 WT and mutants. Anti-HA antibody IPs analysed by western blot.

DHX15 is a prototypic member of the DEAH family of RNA helicases (Jankowsky, 2011). It contains a N-terminal domain of unknown function (residues 1-146), two Rec-A tandem repeats (residues 147-518), WH domain (residues 519-572), Ratched domain (residues 572-671), and OB-fold (residues 671-795) (Fig 2c). The OB-fold is the predominant site of interaction with RNA and proteins, although the Rec-A domains can also establish functional interactions with RNA and protein (Lebaron et al, 2009). Other G-patch containing proteins have been demonstrated to interact with DHX15 via the OB-fold (Chen et al, 2014; Lin et al, 2009; Memet et al, 2017; Niu et al, 2012). To study whether the DHX15 OB-fold is required for CMTR1 binding, N-terminal HA-tagged DHX15 WT or a C-terminal deletion mutant (1-635), ∆Cterm, were transiently expressed in HeLa cells. Endogenous CMTR1 immunoprecipitated with HA-DHX15 WT but not ∆Cterm (Fig 2d).

G-patch domains have the consensus sequence, hhxxxGaxxGxGhGxxxxG, where "G" is glycine, "h" is a bulky, hydrophobic residue (I, L, V, M), "a" is an aromatic residue (F, Y, W), and "x" is any residue (Aravind & Koonin, 1999). The CMTR1 G-patch has this consensus and in addition leucines at residues 94, 106 and 118 which are conserved in the G-patch domains of the established DHX15 interactors, PINX1, NKRF, GPATCH2, RBM5 (Fig 2e) (Chen et al, 2014; Lin et al, 2009; Memet et al, 2017; Niu et al, 2012). Since these conserved leucines in other DHX15-binding proteins are required for the interaction, the impact of mutating CMTR1 L94, L106 and L118 to alanine was investigated using the 2L/A mutant (L94A and L106A), and the 3L/A mutant (L94A, L106A and L118A)(Fig 2e). The 2L/A mutation was sufficient to abrogate the interaction of CMTR1 with DHX15 (Fig 2f). This confirms that CMTR1 interacts with DHX15 through the G-patch domain.

### DHX15 inhibits CMTR1 methyltransferase activity

To interrogate the biochemical function of the CMTR1-DHX15 complex, we investigated the impact of DHX15 on CMTR1 methyltransferase activity. A ^32^P-guanosine-capped transcript was incubated with recombinant CMTR1 and methyl donor, SAM (s-adenosyl homocysteine) (Fig 3a). 1^st^ transcribed nucleotide O-2 methylation (GpppG to GpppGm conversion in our substrate) was observed by thin layer chromatography and quantitated by phosphoimager (Belanger et al, 2010). Incubation of recombinant CMTR1 with equimolar recombinant His^6^-DHX15 resulted in an approximately 50% reduction in O-2 methylation (fig 3a and b). Similarly, His^6^-DHX15 significantly inhibited O-2 methylation of a ^32^P-7-methylguanosine-capped transcript (m7GpppG to m7GpppGm conversion in our substrate) (Fig 3c, d, EV3a, b). His^6^-DHX15 inhibited CMTR1-dependent O-2 methylation in a dose-dependent manner (fig 3c and d). As controls, bovine serum albumin (BSA) did not affect CMTR1-dependent O-2 methylation (Fig EV3c) and ATP did not alter DHX15-dependent repression of CMTR1 (Fig EV3d).

**Figure 3.**
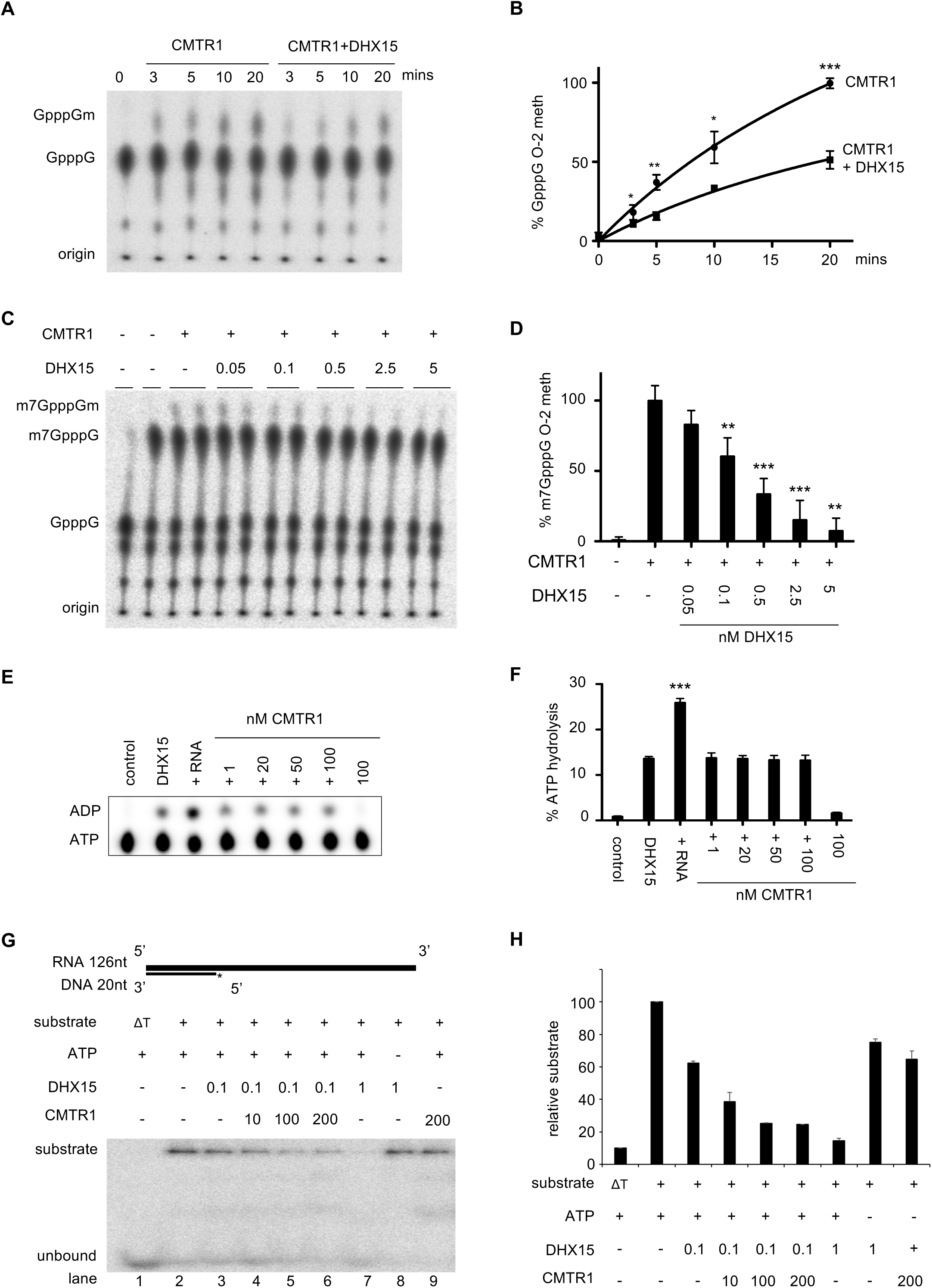
CMTR1 methyltransferase activity is repressed by DHX15. a) GpppG-capped RNA incubated with 3nM CMTR1 and 3nM His^6^-DHX15. Caps throughout figure ^32^P-labelled on a-phosphate. Generation of GpppGm (0-2 methylation) detected by thin layer chromatography (TLC) and autoradiography. b) Percentage conversion of GpppG to GpppGm plotted. Average and standard deviation for three independent experiments. c) m7GpppG-capped RNA incubated with 0.5nM CMTR1 and indicated nM His^6^-DHX15 for 30 mins. Generation of m7GpppGm detected. d) Percentage m7GpppG to m7GpppGm conversion detected. Average and standard deviation of three independent experiments plotted. e) 100nM His^6^-DHX15 incubated with α^32^P-ATP, 1μg total RNA or nM CMTR1 indicated. Generation of ADP detected by TLC and autoradiography. f) Percentage hydrolised ATP. Average and standard deviation depicted for three independent experiments. g) RNA-^32^P-DNA duplex incubated with μM His^6^-DHX15 and nM CMTR1 indicated, and +/-1mM ATP. After incubation for 30min, samples resolved by native PAGE and visualized by phosphorimager. ∆T, 95°C denatured substrate; unbound, single stranded ^32^P-DNA oligonucleotide. h) Average and standard deviation of two independent experiments. Students T test performed relative to b) CMTR1 control, d) CMTR1 alone, f) DHX15 alone. "*", P value<0.05; "**", P value<0.01; "***", P value<0.005.

### CMTR1 increases DHX15 helicase activity

We investigated the impact of CMTR1 on DHX15 ATPase and helicase activity (Figure 3e-h). Established G-patch-containing DHX15 interactors have variable impact on ATPase activity (Lebaron et al, 2009; Tanaka et al, 2007). Recombinant His^6^-DHX15 was incubated with α^32^P-ATP and hydrolysis visualised by the detection of α^32^P-ADP, which was resolved by thin layer chromatography and quantitated by phosphoimager (Fig 3e and f). As established, addition of RNA significantly increased ATP hydrolysis (Fig 3e, f, EV3e, f, g) (Walbott et al, 2010). In order to determine whether CMTR1 influenced DHX15-dependent ATP hydrolysis, a titration of recombinant CMTR1 was included in the ATPase assay. CMTR1 did not activate or repress basal (Fig 3e and f), or RNA-stimulated ATP hydrolysis (Fig EV3h). Addition of SAM had no impact on DHX15 ATP hydrolysis in the presence of CMTR1 or RNA (Fig EV3i).

Since interactors of Prp43, the yeast DHX15 homologue, have been demonstrated to regulate helicase activity independently of ATPase activity (Tanaka et al, 2007), we investigated whether CMTR1 could regulate DHX15 helicase activity. A ^32^P-DNA-RNA duplex was incubated with 1μM His^6^-DHX15. Helicase activity was confirmed by the loss of duplex in the presence but not absence of ATP (Fig 3F, lanes 7 and 8). When 0.1μM His^6^-DHX15 was used in this assay it exhibited minimal helicase activity (lane 3). Addition of CMTR1 increased helicase activity in a dose-dependent manner (lanes 4-6). CMTR1 alone had no helicase activity (lane 9).

### DHX15 prevents CMTR1 recruitment to RNA pol II

CMTR1 interacts with RNA pol II, which is likely to permit efficient co-transcriptional 1^st^ nucleotide O-2 methylation (Haline-Vaz et al, 2008). We investigated the impact of DHX15 on CMTR1 recruitment to RNA pol II. The interaction of recombinant CMTR1 and DHX15 with a biotinylated peptide consisting of three RNA pol II C-terminal domain heptad repeats (YSPTSPS)_3_, unphosphorylated (CTD) or phosphorylated on serine-5 (pCTD) was investigated (Fig4a-c). As a control, RNGTT was demonstrated to interact with pCTD but not CTD (Fig EV4a) (Ho & Shuman, 1999). CMTR1 monomer bound to pCTD but not CTD (Fig 4a), whereas His^6^-DHX15 monomer did not bind to either peptide (Fig 4b). To investigate whether CMTR1 can recruit DHX15 to pCTD, the recombinant proteins were mixed prior to peptide pulldown. Although CMTR1 and DHX15 interact *in vitro* (Fig 1f), CMTR1 was recruited to pCTD whereas DHX15 was not, indicating that in DHX15-CMTR1 complexes DHX15 prevents CMTR1 recruitment to the pCTD (Fig 4c).

**Figure 4.**
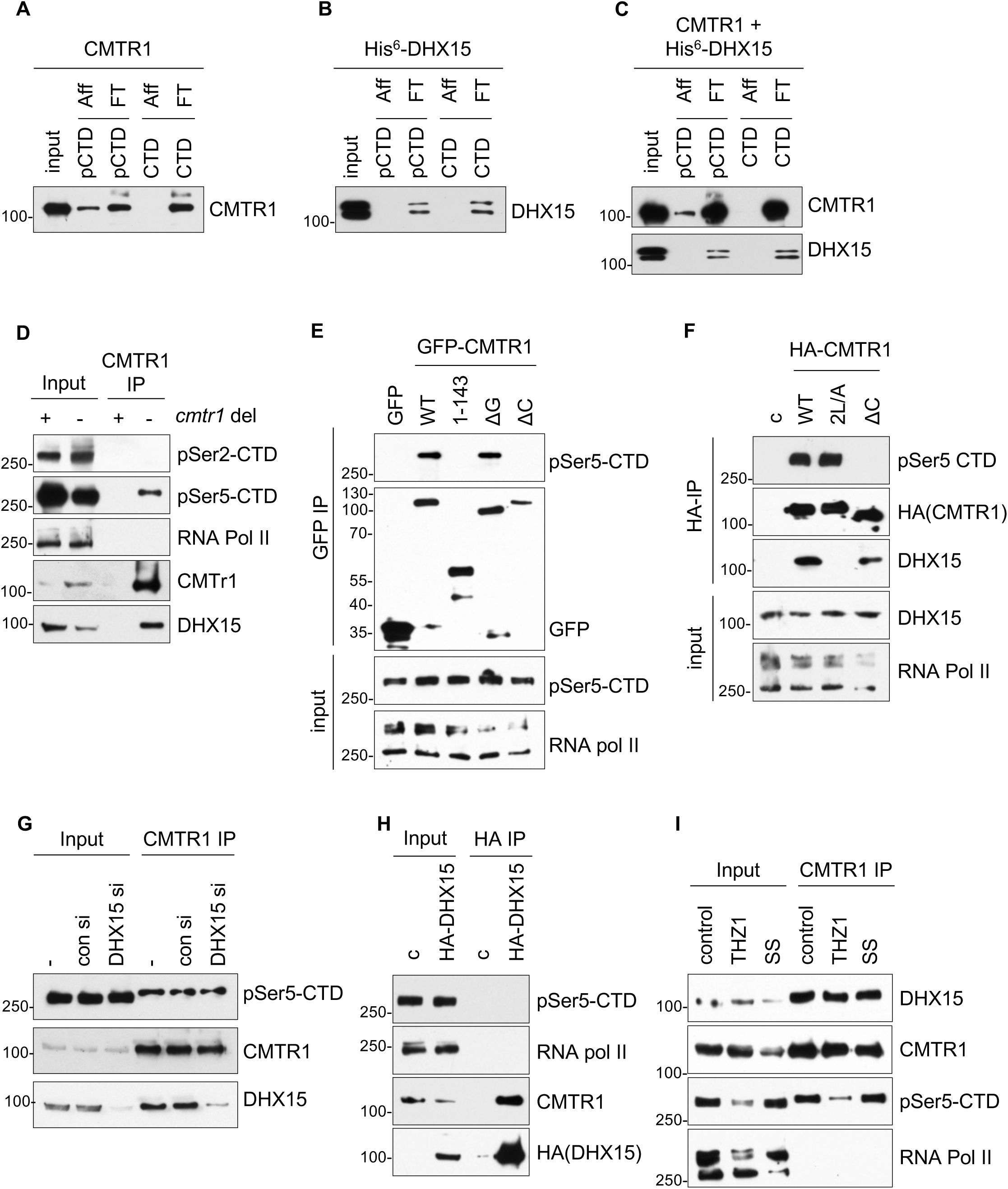
DHX15 inhibits CMTR1 binding to RNA pol II. a) Recombinant CMTR1, b) recombinant His^6^-DHX15, c) pre-mixed CMTR1 and His^6^-DHX15, incubated with biointinylated peptides of 3 RNA pol II CTD heptad repeats, either unphosphorylated (CTD) or phosphorylated on S5 (pCTD). Peptides enriched on streptavidin beads and western blot analysis performed. Aff, affinity purified fraction; FT, flow through. d) CMTR1 IP from HeLa (-) or HeLa *CMTR1* del (+) extracts. Western blot analysis performed. e) pcDNA5 GFP-CMTR1 or indicated mutants expressed in HeLa cells. Anti-GFP antibodies IP analysed by western blot. f) pcDNA 5 HA-CMTR1 or indicated mutants expressed in HeLa cells. Anti-HA antibodies IPs analysed by western blot. g) HeLa cells transfected with DHX15 siRNA or non-targeting control. CMTR1 IPs analysed by western blot. h) pcDNA5 HA-CMTR1 and mutants expressed in HeLa cells. Anti-HA antibodies IPs analysed by western blot. i) HeLa cells treated with 0.5μM THZ1 for 4h or serum starved for 6hrs (SS). CMTR1 IPs analysed by western blot.

The interaction of CMTR1 and DHX15 with RNA pol II was investigated in cells. In HeLa cells extracts, endogenous CMTR1 co-immunoprecipitated with RNA pol II phospho-Ser5-CTD (pSer5-CTD), but not RNA pol II phospho-Ser2-CTD (pSer2-CTD) or unphosphorylated RNA pol II (Fig 4d). As expected, DHX15 was also present in the CMTR1 IPs. The interaction of CMTR1 and pSer5-CTD was RNaseA-insensitive and therefore RNA-independent (FIG EV4b). To map the interaction of CMTR1 with RNA pol II, GFP-CMTR1 WT, 1-143, ∆G and ∆C (1-173), and HA-CMTR1, 2L/A and ∆C were transiently expressed in HeLa cells, immunoprecipitated via their tags and pSer5-CTD binding determined (Fig. 4e and f). GFP-CMTR1 and HA-CMTR1 interacted with pSer5-CTD whereas GFP-CMTR1∆C and HA-CMTR1∆C did not, indicating that the WW domain mediates the interaction with RNA pol II (Fig. 4e and f). GFP-CMTR1∆G and HA-CMTR1 2L/A (both defective for DHX15 binding) interacted with pSer5-CTD, confirming that in cells DHX15 is not required for CMTR1 recruitment to RNA pol II (Fig. 4 e and f). Furthermore, siRNA-mediated suppression of DHX15 expression, did not impact on CMTR1 binding to pSer5-CTD (Fig. 4g).

Although DHX15 is not required for CMTR1 recruitment to pSer5-CTD (Fig 4a, c, e and f), it was important to determine whether in cells DHX15 binds to RNA pol II, either independently of or in a complex with CMTR1. HA-DHX15 was transiently expressed in HeLa cells and anti-HA IP performed (Fig 5h). As expected, HA-DHX15 bound to CMTR1 but not to pSer5-CTD or unphosphorylated RNA pol II, confirming that in cells, as *in vitro,* DHX15 does not appreciably interact with RNA pol II (Fig. 4b, c and h). Furthermore this confirms in cells, as *in vitro,* that CMTR1 in a complex with DHX15 does not interact with RNA pol II (Fig. 5c and h). When the pSer-5-CTD was diminished using THZ1, a CDK7 inhibitor, DHX15 binding to CMTR1 was unaltered, contributing to the evidence that the DHX15-CMTR1 interaction is RNA pol II-independent (Kwiatkowski et al, 2014) (Fig. 4i).

**Figure 5.**
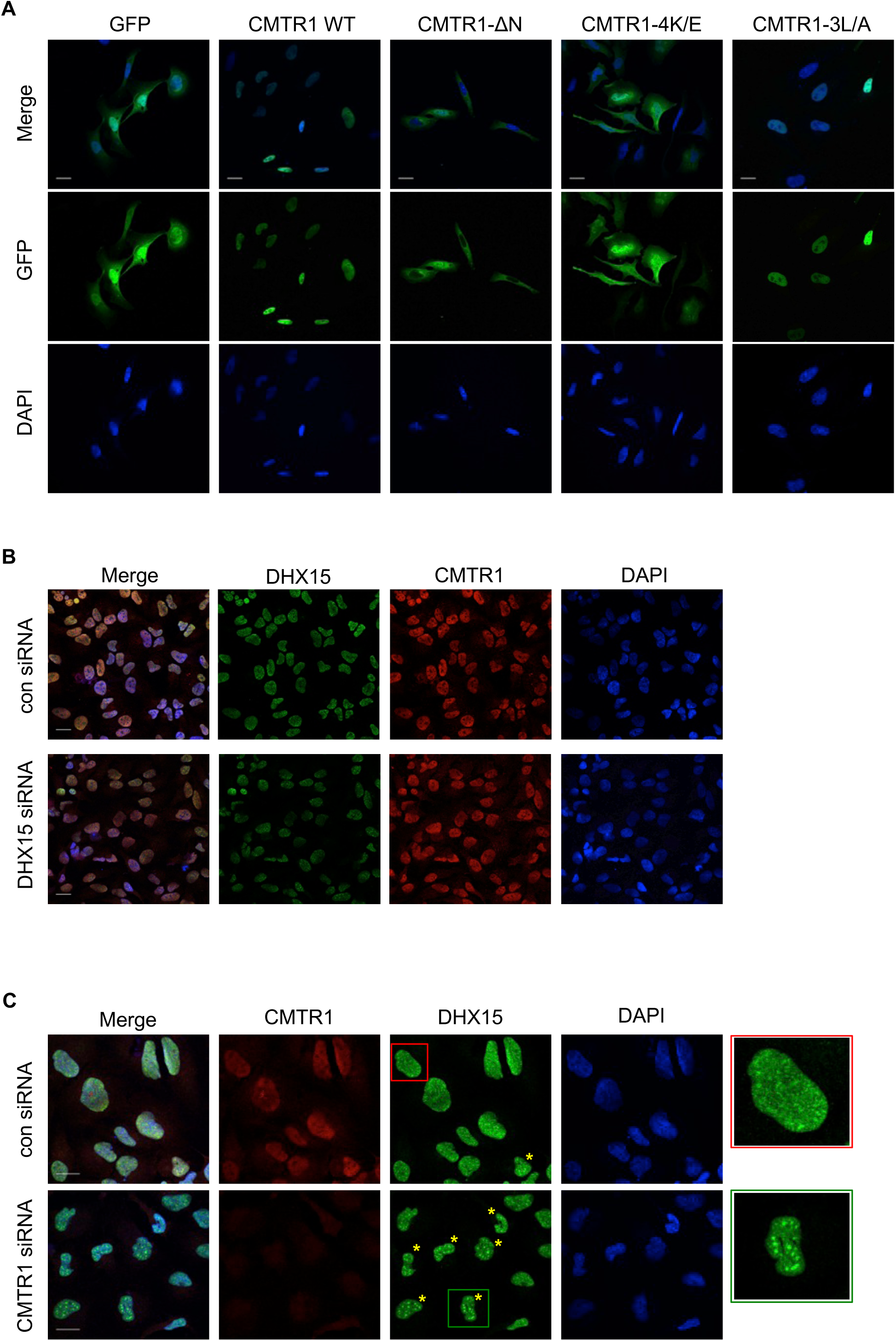
DHX15 localisation is influenced by CMTR1. a) Representative fluorescence images of GFP-CMTR1 or mutants in HeLa cells. b) HeLa cells transfected with DHX15 siRNA or non-targeting control. Representative immunofluorescence (IF) images of endogenous CMTR1 and DHX15. Individual and merged images show CMTR1 (red), DHX15 (green) and DAPI (blue). c) As in (b), except cells transfected with CMTR1 siRNA or control. Yellow asterisks indicate cells with DHX15 accumulation in foci. 4.5x magnified areas marked. Data representative of three independent experiments. Bar indicates 20μm.

### CMTR1 influences DHX15 localisation

Since DHX15 prevents CMTR1 interaction with RNA pol II, we investigated whether DHX15 influences CMTR1 subcellular localisation, which is predominantly nuclear (Haline-Vaz et al, 2008). GFP-CMTR1 and endogenous CMTR1 were predominantly nuclear in HeLa cells (Fig 5a-c). A potential CMTR1 nuclear localisation signal was identified, 14KKQKK (Lange et al, 2007). Mutation of 14KKQKK to 14EEQEE (4K/E) or removal of the first 25 residues (∆N) resulted in partial cytoplasmic localisation, confirming that 14KKQKK contributes to CMTR1 nuclear localisation. In order to determine the impact of DHX15 on CMTR1 localisation, the localisation of GFP-CMTR1 3L/A, which does not bind DHX15, was investigated. GFP-CMTR1 3L/A was predominantly nuclear indicating that DHX15 does not influence CMTR1 localisation (Fig 5a). Furthermore, suppression of DHX15 expression did not alter CMTR1 nuclear localisation (Fig 5b).

Since we had observed that CMTR1 stimulates DHX15 helicase activity *in vitro,* we investigated whether it also influences DHX15 localisation in cells. Prp43, the yeast homologue of DHX15, occupies several cellular locations to execute different biological functions (Heininger et al, 2016). The different Prp43 locations are achieved by the competition of G-patch proteins, which recruit the helicase to different parts of the cell. In HeLa cells, DHX15 exhibited a diffuse nuclear localisation (Fig 5b and c). However, in approximately a third of cells DHX15 also exhibited speckled nuclear foci, a common feature of splicing factors (Tannukit et al, 2009). Indeed, DHX15 foci partially co-localized with the splicing factor SC35 (Pawellek et al, 2014) (Fig EV5a). The number of cell exhibiting DHX15 nuclear foci significantly increased when CMTR1 was knocked-down (Fig 5c and EV5b), or transient transfection of guide RNAs and CRISPR/Cas9 (CMTR1 del cell line, Fig EV5c and d).

### DHX15 represses CMTR1-dependent translation

We had observed that DHX15 inhibits CMTR1-dependent O-2 methylation (Fig. 3), and interaction with RNA pol II (Fig. 4), and that CMTR1 activates DHX15-dependent helicase activity (Fig. 3) and controls DHX15 localisation (Fig. 5). The net outcome of these relationships is complex and will depend on many factors (see discussion). Here, we focussed our investigation on the impact of DHX15 on CMTR1 cellular function using the CMTR1 2L/A mutant, which does not bind to DHX15 (Fig. 2f), but does to RNA pol II (Fig 4f). Since 1^st^ nucleotide O-2 methylation is associated with enhanced translation (Kuge et al, 1998), the impact of the DHX15-CMTR1 interaction on this process was investigated. HCC1806 cells were transfected with pcDNA5 HA-CMTR1 WT, 2L/A or vector control and polysome profiling analysis was performed, in which free ribosomes and ribosomal subunits are separated from translating ribosomes in a sucrose gradient (Fig 6a and b). Although expression of CMTR1 WT and 2L/A had a mild impact on the polysome profiles, across multiple experiments the ratios between polysomes and monosomes were not significantly, indicating that DHX15 was not having a broad impact on translational control (Fig 6c). To investigate gene-specific effects, RNA sequencing analysis was used to quantify total cellular transcripts and the transcripts associated with most dense ribosome binding (Fig 6a, shaded area; Tables 1 and 2). Out of 12238 gene transcripts that passed quality thresholds, none exhibited a significant difference in expression level in cells expressing CMTR1 WT and 2L/A (Table 1, Figure 6d). This indicates that in HCC1806 cells, the CMTR1-DHX15 interaction is unlikely to have a significant impact on transcription or RNA stability. However, 59 genes were significantly enriched in polysomes in cells expressing CMTR1 2L/A compared to WT, indicating that the DHX15-CMTR1 interaction restricts the translation of a subset of mRNAs (Fig. 6e and f; table 2). Conversely, no genes were significantly depleted from polysomes in cells expressing CMTR1 2L/A compared to WT, indicating that the DHX15-CMTR1 interaction does not enhance the translation of any mRNA.

**Figure 6.**
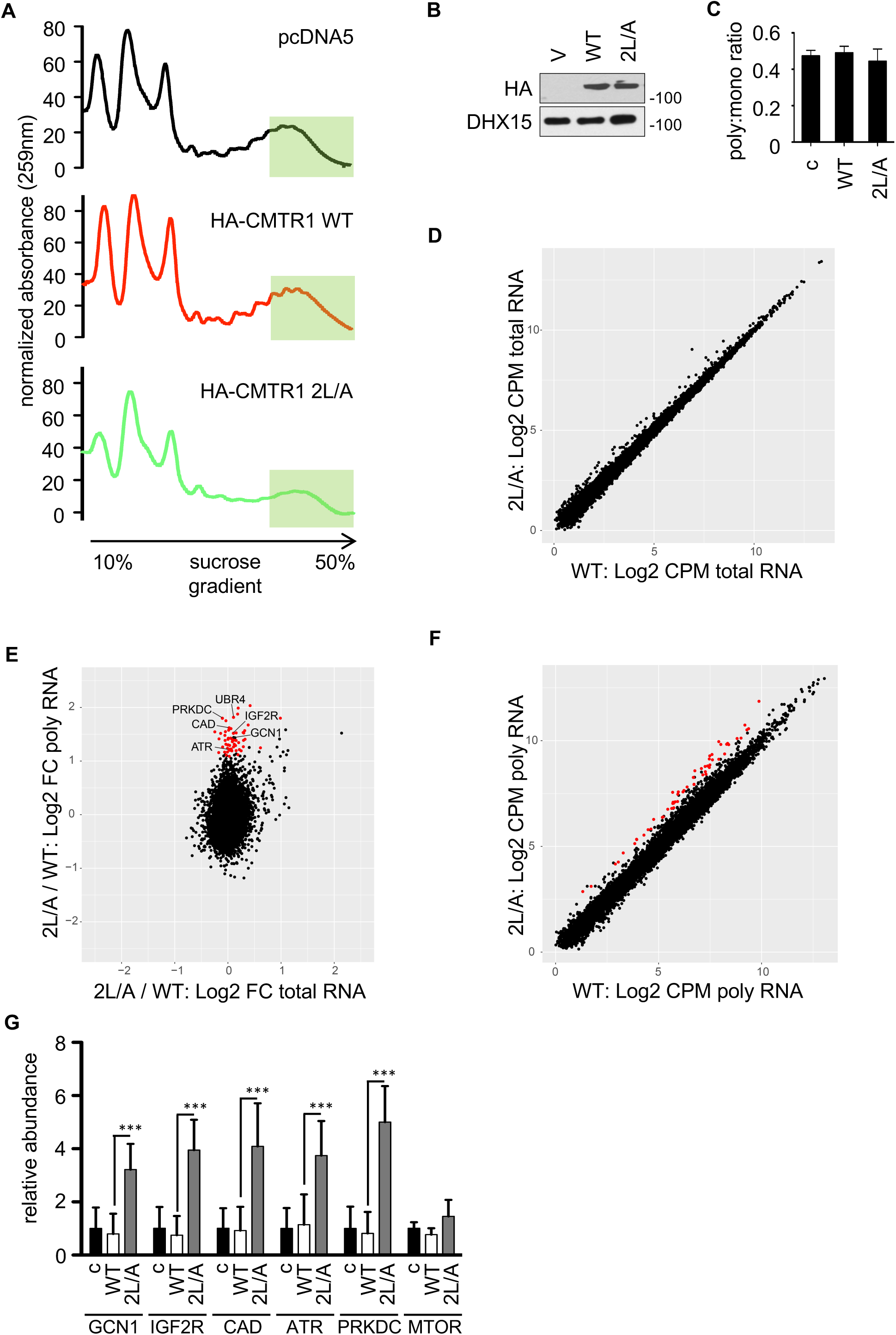
The DHX15-CMTR1 interaction inhibits translation of a subset of genes. a) Extracts from HCC1806 cells transfected with indicated constructs resolved on 10-50% sucrose gradients. Representative 259nm absorbance for 4 independent experiments. Fraction analysed by RNA seq indicated by shaded box. b) Cell extracts analysed by western blot. c) Average and standard deviation of ratio of polysome to mononosome absorbance for 4 independent experiments. RNAseq analysis used to determine uniquely aligned reads per gene (transcript isoforms collapsed) in RNA samples from HCC1806 cells expressing HA-CMTR1 WT and 2L/A (n=2). d) Scatter plot of Log2 transformed total counts per million (Log2CPM) in total RNA. e) Scatter plot of log2 fold change (FC) in 2L/A / WT for total RNA and polysomal RNA. Significantly regulated genes determined by a negative binomial rest in EdgeR in red. f) Scatter plot of Log2 transformed counts per million (Log2CPM) in polysomal RNA. g) Polysomal mRNA levels quantified by RTPCR. Average and standard deviation for three independent experiments. Students T test performed "***", P value<0.005.

The genes which exhibited enhanced translation in cells expressing CMTR1 2L/A were enriched for GO terms associated with important metabolic functions and cell cycle processes (Table 3). Notably there was enrichment for genes encoding factors involved with the cell cycle and DNA damage responses including ATR, UBR4, UBR2, CENPE, PRKDC and BIRC6; factors involved in RNA processing including GCN1, TPR, RANBP2 and NUP205; metabolic enzymes including FASN, EPRS, and CAD; and focal adhesion-associated molecules HSPG2, FLNB, MYH9, FAT1, IGF2R, and CLTC. The polysome loading of selected genes was confirmed by RTPCR (Fig. 6g). Taken together this analysis indicates that DHX15 may repress the positive effect of CMTR1 on cell growth and proliferation.

### The interaction of DHX15 and CMTR1 inhibits proliferation

The impact of CMTR1 expression on cell proliferation was investigated in HCC1806 cells and another mammary epithelial tumour line, MCF7. Transfection of two independent CMTR1 siRNAs resulted in suppression of CMTR1 expression and a reduction in cell proliferation (Fig. EV1b, 7a-c). Given that expression of CMTR1 2L/A resulted increased polysome loading of genes involved in metabolism and cell cycle, we investigated its impact on cell proliferation. When HA-CMTR1 was transiently expressed in sparsely plated HCC1806 cells, a significant increase in cell number was observed after 24 hours (Fig. 7d). Transient expression of HA-CMTR1 2L/A resulted in a further increase in cell number, supporting the hypothesis that DHX15 suppresses the translation of a subset of pro-growth genes. To investigate if the increased expression of these genes may contribute to the enhanced proliferation observed on expression of CMTR1 2L/A, two genes with increased polysome loading in CMTR1 2L/A cells, CAD and GCN1, were suppressed by siRNA transfection (Fig. 7e). Suppression of CAD and GCN1 resulted in reduced proliferation of HCC1806 cells, supporting the hypothesis that DHX15 controls CMTR1-dependent translation of a subset of pro-growth genes.

**Figure 7.**
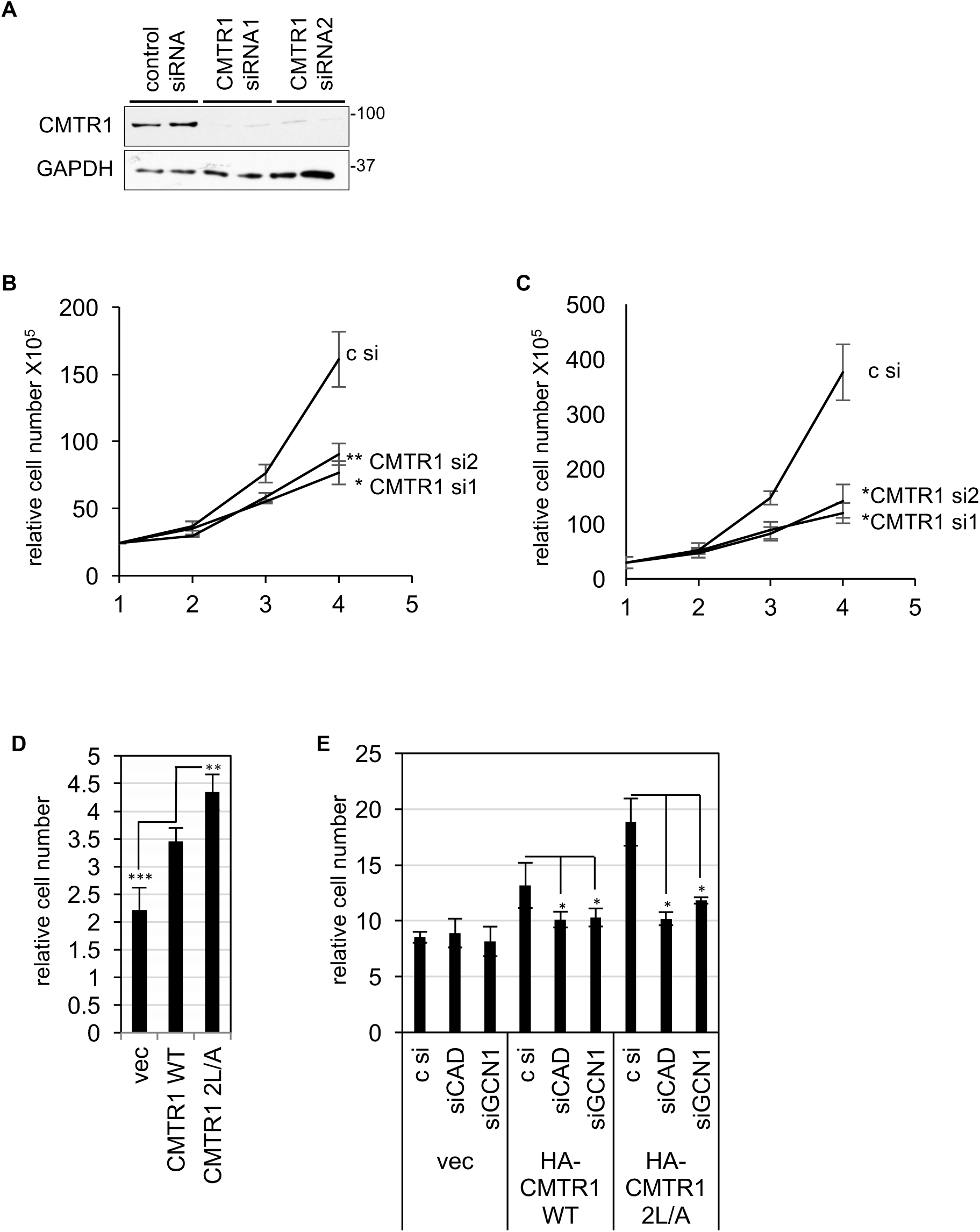
The DHX15-CMTR1 interaction inhibits cell proliferation. a) MCF7 cells transfected with 2 independent siRNAs or non-targeting control. Cell extracts analysed by western blot. b) MCF7, c) HCC1806 cells transfected with 2 independent siRNAs or non-targeting control. Cells counted every day. HCC1806 transfected with d) pcDNA5 CMTR1 WT or 2L/A or vector control, e) the same plasmids and 50nM CAD, GCN1 or non-targeting control siRNA. After 48 hours cells counted. Average and standard deviation for 3-5 independent wells. Students T test performed. "*", P value<0.05; "**", P value<0.01; "***", P value<0.005.

## Discussion

Higher eukaryotes carry mRNA cap modifications not present in lower eukaryotes including O-2 methylation of the 1^st^ transcribed nucleotide ribose, which is associated with translation and the identification of self RNA in innate immunity (Kuge et al, 1998; Leung & Amarasinghe, 2016; Schuberth-Wagner et al, 2015). Increased complexity of the mRNA capping enzymes is also observed in higher eukaryotes, including additional domains and subunits (Aregger & Cowling, 2013; Gonatopoulos-Pournatzis et al, 2011; Ramanathan et al, 2016; Shuman, 2002). The additional complexity of the mammalian capping enzymes facilitates regulation of mRNA cap formation by cellular signaling pathways, which impacts on gene regulation and thus cell physiology and function (Aregger et al, 2016; Grasso et al, 2016).

We investigated mechanisms regulating CMTR1, the 1^st^ nucleotide ribose O-2 methyltransferase (Belanger et al, 2010). We isolated CMTR1 from HeLa cells as a monomer and in a complex with DHX15, a DEAH-box helicase involved in RNA processing, including splicing and ribosome biogenesis (Koodathingal & Staley, 2013; Robert-Paganin et al, 2015). DHX15 lacks RNA-sequence specificity, but possess a C-terminal OB-fold through which it binds to a series of G-patch-containing proteins, allowing the helicase to contribute to several nuclear functions (Chen et al, 2014; Memet et al, 2017; Niu et al, 2012; Robert-Paganin et al, 2015; Tauchert et al, 2017). CMTR1 contains a G-patch and we identified DHX15 as the predominant CMTR1-interacting protein. Mutating two conserved leucines in the CMTR1 G-patch (2L/A mutant), was sufficient to abrogate interaction with DHX15. Previous studies had suggested that CMTR1 and DHX15 participate in the same mRNA processing events (Gebhardt et al, 2015; Yoshimoto et al, 2014). Here we demonstrate that the direct interaction of CMTR1 and DHX15 regulates the catalytic activities and localisation of both enzymes, which impacts on gene expression and cell proliferation.

### DHX15 restrains CMTR1 function

mRNA cap formation initiates during the early stages of transcription, when the RNA triphosphatase/guanylyltransferase (RNGTT) and the N-7 cap guanosine methyltransferase (RNMT) are recruited to Ser-5 phosphorylated RNA pol II CTD (Ramanathan et al, 2016). We report that CMTR1 interacts directly with Ser-5 phosphorylated RNA pol II CTD, which is likely to facilitate efficient co-transcriptional 1^st^ nucleotide O-2 methylation. DHX15 inhibits recruitment of CMTR1 to RNA pol II and equimolar DHX15 reduces CMTR1 methyltransferase activity by half. Thus the interaction of DHX15 with CMTR1 is likely to constrain 1^st^ nucleotide O-2 methylation to the initial stages of transcription, and suppress or restrain post-transcriptional, aberrant methylation. To our knowledge, this is the first example of DHX15 regulating the catalytic activity of an interacting protein. Furthermore, CMTR1 is the only annotated protein containing both a G-patch domain and a catalytic domain.

Previous studies demonstrated that O-2 methylation is associated with translation efficiency and promotes translation of oocyte maternal mRNAs (Kuge et al, 1998; Kuge & Richter, 1995; Muthukrishnan et al, 1976a). In order to investigate the impact of DHX15 on CMTR1-dependent translation, the CMTR1 2L/A mutant (which does not bind to DHX15) was expressed, resulting in increased ribosome loading of a subset of transcripts. Therefore DHX15 restricts the translation of a specific subset of genes, which include those involved in metabolic pathways and cell cycle control. Expression of CMTR1 2L/A also resulted in increased cell proliferation.

The gene-specific impact of the DHX15-CMTR1 interaction parallels observations made following regulation of N-7 cap guanosine methylation; CDK1-dependent regulation of RNMT activity and ERK-dependent regulation of RAM expression both result in specific changes in gene expression, which influence cell proliferation and embryonic stem cell differentiation, respectively (Aregger et al, 2016; Grasso et al, 2016). The mechanisms responsible for these gene-specific effects are currently unclear. The genes sensitive to the DHX15-CMTR1 interaction may be particularly dependent on O-2 methylation for polysome loading or these genes may require high levels of CMTR1 activity to be O-2 methylated.

### CMTR1 controls DHX15 function

The mechanism by which G-patch proteins regulate DHX15 activity and localisation has been characterised for several other interactors. DHX15 uses energy generated by ATP hydrolysis to power helicase activity (Walbott et al, 2010). The DHX15 RecA domains interact with the C-terminal domain, which renders the enzyme in a closed conformation. Disruption of these interactions by ATP binding promotes changes in the helicase structure, leading to the open conformation required for unwinding complex RNA stretches (Tauchert et al, 2017). In addition, the stacking of the adenosine base with the RecA1 R- and RecA2 F-motifs is important for the activity and regulation of the helicase (Robert-Paganin et al, 2017). When intrinsically unstructured G-patch domains bind to the DHX15 OB-fold, they can adopt a stable secondary structure (Christian et al, 2014). G-patch domains can influence both the stacking of the adenosine base and interactions of N- and C-terminal domains, thus regulating DHX15 catalytic activity (Robert-Paganin et al, 2017; Tauchert et al, 2017). In accordance, we observed that CMTR1 activates DHX15 helicase activity. However, although several G-patch proteins increase ATPase activity, CMTR1 did not affect basal or RNA-stimulated DHX15 ATPase activity (Robert-Paganin et al, 2017). A similar observation was made with the yeast splicing factor Ntr1, which did not alter Prp43 ATPase activity but did activate the helicase (Tanaka et al, 2007). Furthermore, although the G-patch is defined by well-conserved residues, there are now several examples of G-patch proteins impacting differentially on DHX15 activity and function (Aravind & Koonin, 1999; Banerjee et al, 2015; Tauchert et al, 2017).

The competition of cofactors for DHX15 was previously observed to regulate its distribution between different pathways (Heininger et al, 2016). DHX15 interacts with splicing factors, including TFIP11, and localises to nuclear splicing speckles (Tannukit et al, 2009). We also detected DHX15 co-localised with the splicing factor SC35 in nuclear speckles. Suppression of CMTR1 expression resulted in increased occurrence of DHX15 in nuclear speckles, indicating that CMTR1 also competes with other G-patch proteins for DHX15 binding.

### Co-ordinated impact of DHX15 and CMTR1 on gene expression

Ultimately we want to understand how the relationship between DHX15 and CMTR1 impacts on cell physiology and function, which is complex due to the multifunctional nature of both enzymes. The cellular impact of the DHX15-CMTR1 interaction will depend on the relative expression of the enzymes and that of their other interacting partners, which also influence enzyme activity and localisation (e.g. pSer5-RNA pol II for CMTR1 and RNA processing factors for DHX15). The impact of the DHX15-CMTR1 interaction will also depend on the physiology of the cell, including the relative dependency on O-2 methylation, splicing, rRNA processing and translation. Therefore it is likely that the net outcome of the DHX15 and CMTR1 interaction will vary in different cell lineages and under different physiological circumstances.

Recently, somatic mutations of DHX15 were shown to be recurrent in the development of Core-Binding Factor Acute Myeloid Leukemia (CBF-AML). In particular, the missense DHX15-Arg 222 Gly mutation is present in ~5% of CBF-AML patients, both at diagnosis and/or disease relapse (Faber et al, 2016; Farrar et al, 2016; Sood et al, 2016). DHX15-Arg222 is part of the RecA1-β-Turn RF conserved motif (Arginine-Phenylalanine), which is a key player in RNA translocation (Tauchert et al, 2017). Interestingly, DHX15-R222G has reduced interaction with proteins involved in splicing (TFP11), ribosomal biogenesis (NFRP) and translation (CMTR1), which may contribute the impact of the DHX15 mutant.

In summary, CMTR1 is regulated by DHX15 impacting on gene expression and cell physiology. We now recognise that the mRNA capping enzymes, RNGTT, RNMT and CMTR1 are regulated by different co-factors and posttranslational modifications, with multiple impacts on gene expression and cell physiology. The challenge going forward is to understand the complex integration of these regulatory events during development and in the adult.

## Materials and Methods

### Cell Culture

HeLa, HEK293, MCF7 and U2OS cells cultured in DMEM and HCC1806 cells cultured in RPMI, both with 10% (v/v) Foetal Bovine Serum, 100U/ml penicillin, 0.1mg/ml streptomycin and 2mM L-glutamine. For plasmid transfection, cells in 10 cm dishes transfected with 1-5μg plasmid using 1 mg/ml polyethylenimine (Polysciences). Cells cultured for 36 hours before lysis in 0.45ml ice-cold lysis buffer (50mM Tris/HCl pH 7.5, 1mM EGTA, 1mM EDTA, 1mM sodium orthovanadate, 10mM β-glycerophosphate, 50mM NaF, 5mM sodium pyrophosphate, 0.27M sucrose) supplemented with 1% (v/v) Triton X-100, 2mM DTT, 1% (v/v) Apoprotin, 10μM Leupeptin and 1μM Pepstatin. Extracts centrifuged at 16200 g at 4°C for 15 mins and supernatant retained. For siRNA transfections, cells transfected with 50nM siRNA (Dharmacon siGenome range; non-targeting or CMTR1), for 48 hours unless stated otherwise. UAGAUGAUGUUCGGGAUUA, 01 CMTR1 siRNA; GUAAGAGCGUGUUUGAUGU, 02 CMTR1 siRNA. For plasmid and siRNA co-transfection, lipofectamine 2000 used. 2-6 x 10^4^ cells plated in 6 well plate, 0.5μg pcDNA5 GFP and 0.5μg pcDNA5 HA-CMTR1 WT or 2L/A mixed with siRNA prior to transfection. Countess cell counter was used for cell counting.

### Generation of a CMTR1 knockout cell line (CMTR1 del) using CRISPR/Cas9

CMTR1 gene (ensembl ENSG00000137200) optimal scoring guide target site GGGAGGTTCATCATCGGACG[TGG] cloned into pBabeD pU6 and sequence-verified (http://crispr.mit.edu/). HeLa cells stably expressing Cas9 co-transfected with 3 μg pBabeD pU6 CMTR1 using Lipofectamine 2000. Cells incubated in DMEM, 10% fetal bovine serum (FBS), 2 mM L-glutamine, 100 μg/ml NormocinTM (Invivogene). After 12 hrs, medium replaced by fresh medium /4 μg/ml Puromycin. After 48 hrs selection, 2μg/ml doxycycline added to induce Cas9 expression. After 72hrs, single clones FACS-sorted into 96 well plate containing DMEM/20% fetal bovine serum (FBS), 2 mM L-glutamine, 100 units/ml penicillin, 100 μg/ml streptomycin and 100 μg/ml NormocinTM (Invivogene). CMTR1 protein expression was screened by western blot in 40 clones. For CMTR1 del clones displaying absence of protein, genomic DNA isolated and amplified by PCR, using CMTR1 genomic forward: CTGTGATCCCAGTGGCTGT and CMTR1 genomic reverse: CCAAGGGGCAGTGGACTAT primers. PCR product from control and CMTR1 del clones sequenced. CMTR1 del clone 5 had no region homology to WT cells around Cas9 targeting sequence and selected for the experiments.

### Immunoprecipitation and Western Blot

For immunoprecipitation (IP) of GFP or HA-tagged proteins, anti-GFP antibody conjugated agarose (Chromotek) and anti-HA antibody conjugated agarose (Sigma) used. For endogenous proteins, 1μg relevant antibody pre-incubated with 10μ1 protein-G Sepharose packed beads and washed to remove non-bound antibody. 0.5-2.5mg lysates precleared with 10-30μ1 protein-G Sepharose (GE Healthcare), then incubated with antibody-resin conjugate for 2 hrs at 4°C under gentle agitation. IPs washed three times with lysis buffer containing 0.1M NaCl. RNAseA treatment: 40μg incubated with IP or 2μg HeLa RNA for 60 min at 4°C. Proteins eluted in SDS-sample buffer. Western blots performed by standard protocols. CMTR1 antibody raised in sheep by Orygen Antibodies Limited, UK and affinity purified against the human recombinant protein. 2^nd^ bleed used at 1μg/ml for western blotting and the 1^st^ bleed used at 1μg/IP. Other antibodies: RNGTT (in-house), HA (Sigma), DHX15 (Abcam, ab70454), GAPDH (Cell Signaling Technologies), RNA pol-II (N20, Santa Cruz), PSer5-PolII (3E8, Chromotek) and pSer2-PolII (3E10, Chromotek). Secondary antibodies from Pierce.

### Mass Spectrometry analysis

IPs resolved by SDS-PAGE and stained with Novex Colloidal blue (Invitrogen). Bands excised and washed sequentially on vibrating platform with 0.5ml water, 1:1(v/v) water and acetonitrile (AcN), 0.1M ammonium bicarbonate, 1:1(v/v) 0.1M ammonium bicarbonate and AcN. Samples reduced in 10mM dithiothreitol (20 min, 37°C), alkylated in 50mM iodoacetamide/0.1M ammonium bicarbonate (20 min, dark), and washed in 50mM ammonium bicarbonate and 50mM ammonium bicarbonate/50% AcN. When gel pieces colourless they were washed with AcN for 15min and dried. Gel pieces incubated (16h, 30°C) in 5μg/ml trypsin/25mM triethylammonium bicarbonate. Tryptic digests analysed by liquid chromatography–mass spectrometry on Applied Biosystems 4000 QTRAP system (Foster City, CA) with precursor ion scanning. Resulting MS/MS data searched using the Mascot search algorithm (http://www.matrixscience.com).

### Bacterial Expression and Purification of DHX15 and CMTR1 proteins

Bacterial expression plasmids pET15b-His^6^-DHX15 or pGEX6P1-GST-3C-CMTR1 transformed into Rosetta 2 (DE3) cells (Novagen) and plated on LB agar/100ug/ml ampicillin and 35ug/ml chloramphenicol. Single colonies picked to inoculate LB starter cultures with above antibiotics and incubated at 37°C overnight. After 16 hrs, cultures diluted 1/50 into fresh 6-9L LB cultures and incubated at 37°C with agitation. At OD600 ~ 0.3-0.4, temperature reduced to 16°C, and at OD600 ~ 0.7-0.8 protein expression induced with 50uM IPTG for 14-16hrs. Cultures harvested by centrifugation and pellets resuspended in ice-cold re-suspension buffer (Tris 50mM pH 8, 200mM NaCl, 1mM DTT, 25mM imidazole, 10% glycerol (v/v) and Complete-EDTA free protease inhibitor from Roche). Resuspended cells flash-frozen in liquid nitrogen, thawed in a lukewarm (25°C) water bath and treated with lysozyme (1mg/mL) and DNase I (1:10000 dilution) for 30min at 4°C. Cell debris and insoluble material clarified by centrifugation at 38000 RCF at 4°C for 45min. Soluble lysates passed through poly-prep columns packed with either Ni-NTA agarose (His^6^-DHX15) or GSH-sepharose (GST-3C-CMTR1) pre-equilibrated in wash buffer (50mM Tris pH 8, 500mM NaCl, 1mM DTT, 25mM imidazole, 5% glycerol (v/v)). Post-binding, resins were extensively washed with wash buffer followed by a final wash step in low salt (200mM NaCl) wash buffer. For His^6^-DHX15, Ni-NTA bound material eluted with elution buffer (50mM Tris pH 8, 200mM NaCl, 1mM DTT, 5% glycerol and 300mM imidazole, while GSH-sepharose bound GST-CMTR1 material was treated with GST-3C protease overnight at 4°C to cleave GST tag. Eluted His^6^-DHX15 and cleaved CMTR1 proteins concentrated using Amicon 30 kDa-MWCO concentrator and subject to size-exclusion chromatography (Superdex 200) in 50mM Tris pH 8, 200mM NaCl, 1mM DTT and 5% glycerol (v/v) buffer on an AKTA purifier FPLC system. Proteins further concentrated and flash-frozen as single use aliquots, stored at - 80°C.

### Gel filtration

0.5ml 2mg/ml cell extract 0.22-μm filtered and resolved on Superdex 200 10/300 GL preparative grade column (GE Healthcare) in 50mM Tris–HCl pH 7.4, 1mM EDTA, 0.1M sodium chloride, 0.03% Brij-35 and 2mM DTT at 0.4ml/min flow. 0.5ml fractions collected. Molecular weight markers (Bio-Rad) were used: bovine thyroglobulin (670kDa), bovine gamma globulin (158kDa) and chicken ovalbumin (44kDa).

### First nucleotide O-2 methylation assay

A guanosine-capped substrate ^32^P-labelled on α-phosphate (GpppG-RNA) was produced. 200 ng 55-base *in vitro*-transcribed RNA incubated with 100 ng RNGTT, 2 μl (10 μCi) [α-^32^P]GTP and 1 μl RNAsin (Promega) in 10 μl reaction buffer (0.05 M Tris/HCl, pH 8.0, 6 mM KCl and 1.25 mM MgCl_2_) at 37°C for 60 mins. When used, guanosine-capped transcript incubated with 100ng RNMT and 0.2 μM s-adenosyl methionine (SAM) for 20 min at 30°C to produce N-7 methylated guanosine cap (m7GpppG-RNA). RNA purified by ammonium acetate precipitation. In methyltransferase assay, indicated concentration of CMTR1 incubated with ^32^P-labeled GpppG-RNA or m7GpppG-RNA and 10μM SAM at 30°C for 30 min (unless otherwise stated). Following reaction, RNA precipitated and resuspended in 4 μl 50 mM sodium acetate and 0.25 units P1 nuclease for 30 min at 37°C. Cap structures resolved in 0.4 M ammonium sulphate on polyethyleneimine–cellulose plates, and visualized and quantified by autoradiography.

### Molecular biology

cDNA cloning and mutagenesis performed by standard protocols. Constructs sequence-verified.

### CTD peptide affinity chromatography

RNA pol-II CTD peptide chromatography performed as in (Ho & Shuman, 1999). 0.5nmol biotinylated CTD peptides absorbed on 0.2mg Streptavidin Dynabeads M-280 (Invitrogen) in 300μl Binding Buffer (25mM Tris HCl, pH 8, 50mM NaCl, 1mM DTT, 5% Glycerol, 0.03% Triton X-100). After 45min at 4°C, beads magnet-concentrated and washed in binding buffer. 0.2μg indicated protein mixed with peptide-beads in 50μl binding buffer. After incubation for 45 mins at 4°C, beads washed three times with binding buffer and bound fraction eluted with 30μl SDS-PAGE loading buffer and at 95°C for 5mins.

### ATPase activity assay

ATPase reactions performed as in (Lebaron et al, 2009). 100nM DHX15 in 5μl 25-mM Tris-acetate (pH 8.0), 2.5mM Mg(CH3-COO)_2_, 100mM KCl, 0.2mM DTT, 100μg/ml bovine serum albumin (Sigma), 0.6μCi/μl [alpha-32P] ATP and 100-200 μM ATP incubated at 30°C for time indicated. 1μl resolved on polyethyleneimine-cellulose plates using 0.75M KH_2_PO_4_. Spots visualised and quantified on Phosphorimager. When indicated, 1μg HeLa cell RNA added to reaction.

### Unwinding Assay

Unwinding assays performed as in (Tanaka et al, 2007). The 126-nucleotide (nt) RNA strand (5' GGGCGAAUUGGGCCCGACGUCGCAUGCUCCCGGCCGCCAUGGCGGCCGCGGGAAUUCGAUUAUCACUAGUGAAUUCGCGGCCGCCUGCAGGUCGACCAUAUGGGAGAGCUCCCAACGCGUUGGAUG 3') was synthesized by in vitro transcription, and annealed at a 1:3 molar ratio to 5' ^32^P-labelled DNA (5^’^ GACGTCGGGCCCAATTCGCCC 3’) to yield 5'-tailed 30-bp duplex. Annealed substrate gel-purified on native 6% PAGE. 10 μl reaction mixtures containing 40 mM Tris-HCl (pH 7.0), 2 mM DTT, 1 mM ATP-Mg^2^+, 2.5 nM RNA/DNA substrate, and indicated concentrations of His^6^-DHX15 and CMTR1 incubated at 37°C for 45min. Reactions halted by transfer to ice and addition of 5-μl l loading buffer (100 mM Tris-HCl at pH 7.4, 5 mM EDTA, 0.5% SDS, 50% glycerol, 0.1% [w/v] bromophenol blue, xylene cyanol dyes). Samples resolved on 16% polyacrylamide gel in 40 mM Tris-borate, 0.5 mM EDTA, and 0.1% SDS. ^32^P-labeled substrate and products visualized by autoradiography.

### Immunofluorescence

Incubations performed in 2%BSA/PBS at room temperature unless stated. Cells fixed in 4% paraformaldehyde for 10 min, permeabilized with 1% NP-40 / PBS for 3 min, blocked with 10% donkey serum for 20 min, and incubated in 1.4 μg/ml polyclonal sheep anti-CMTR1 or DHX15 antibodies for 1.5 hr, then washed and incubated with 1.4 μg/ml Alexa Fluor 594 or 488-conjugated Donkey Anti-Sheep or Anti-Rabbit antibodies for 45 min. Cells counterstained with 1 μg/ml DAPI (4',6-diamidino-2-phenylindole), mounted in 2.5% DABCO, and visualized by fluorescence microscopy (Zeiss LSM 700).

### Polysome profile

Cells incubated in 100 μg/ml cycloheximide for 10 min, washed in ice cold PBS supplemented with 100 μg/ml cycloheximide and extracts collected in polysome lysis buffer (15mM Tris (pH7.5), 15mM MgCl_2_, 0.3 M NaCl, 1 mM DTT, 1% Triton X-100, 100 mg/ml cycloheximide, 100 U/ml RNasin). 10% extracts retained as input and 90% resolved by centrifugation through 10 ml 10%–50% sucrose gradient at 18 000 x g for 2 hr at 4°C. Fractions collected on FoxyR1 fractionator (TELEDINE ISCO) with OD259 nm monitoring.

### RNA-sequencing and analysis

Extraction of polysomal RNA performed as described previously (Grasso et al, 2016). Briefly, polysomal fractions 16-20 purified by Phenol:Chloroform:Isoamyl Alcohol (25:24:1). RNA precipitated overnight with 2M of LiCl, 20mM Tris pH 7.5, EDTA 10mM. Input RNA purified using Trizol Reagent. RNA sequenced at Tayside Centre for Genomic Analysis. RNA quality checked using TapeStation (Agilent Technologies). RNA sequencing libraries prepared with TruSeq Stranded Total RNA with Ribo-Zero-Gold kit (Illumina). Sequencing performed using NextSeq Series High Output Kit 2 × 75 bp (Illumina). Reads quality controlled using FastQC, then mapped to (GRCh38/hg25) assembly of human genome and reads per gene quantified using STAR 2.5.2b (Dobin et al., 2013). Differentially expressed genes identified using edgeR package (Robinson et al., 2010). Genes with at least 1 count per million (CPM) in all samples analysed for differential expression. Pairwise comparisons made between input RNA and between polysomal RNA for samples transfected with pcDNA5, pcDNA 5 HA-CMTR1 WT or 2L/A. Plots comparing differential input or polysome mRNA abundance drawn using the ggplot2 R package.

### RTPCR

RNA extracted using Trizol Reagent (Invitrogen). cDNA synthesised using iScript cDNA Synthesis Kit (Bio-Rad). RT-PCR performed using Quanta Biosciences SYBR Green. Primers: PRKDC_Fwd GAGAAGGCGGCTTACCTGAG, PRKDC_Rvr CGAAGGCCCGCTTTAAGAGA, IGF2R_Fwd AGCGAGAGCCAAGTGAACTC, IGF2R_Rvr TCGCTGTAAGCAGCTGTGAA, CAD_Fwd AGGTTTGCCAGCTGAGGAG, CAD_Rvr TAATGAGTGCAGCAGGGGTG, ATR_Fwd GGAGGAGTTTTGGCCTCCAC, ATR_Rvr TGTGGCACTGCCCAGCTC, GCN1_Fwd CTTGTGCCCAAGCTGACAAC, GCN1_Rvr GCCCTGTGTCATCCTCTACG, MTOR_Fwd GAAGCCGCGCGAACCT, MTOR_Rvr CTGGTTTCCTCATTCCGGCT, CMTR1_Fwd CATTGCCCCATTTCACATTTGC, CMTR1_Rvr TCTTAGGCCCTGTGCATCTG

## Acknowledgements

We thank the Cowling lab for assistance and Janusz Bujnicki lab, Frances Fuller-Pace and Sara Ten-Have for discussions. Research funded by MRC Senior Fellowship (VC), Lister Institute Prize Fellowship (VC), Wellcome Trust Centre award 097945/Z/11/Z and the Dundee Imaging Facility (Wellcome Trust Technology Platform award 097945/B/11/Z; MRC Next Generation Optical Microscopy award MR/K015869/1), the Division of Signal Transduction Therapy: Boehringer-Ingelheim, GSK, Merck KgaA. THZ-1 was a gift from Nathanael Gray (Dana Farber Cancer Institute).

## Author Contributions

FIV and VHC designed and performed experiments, and wrote the manuscript. AG, LC, ARF, VHC, SW and MP made reagents and performed experiments.

## Conflict of Interest

Authors have no conflicts of interest to disclose.

---

**Figure EV1.**
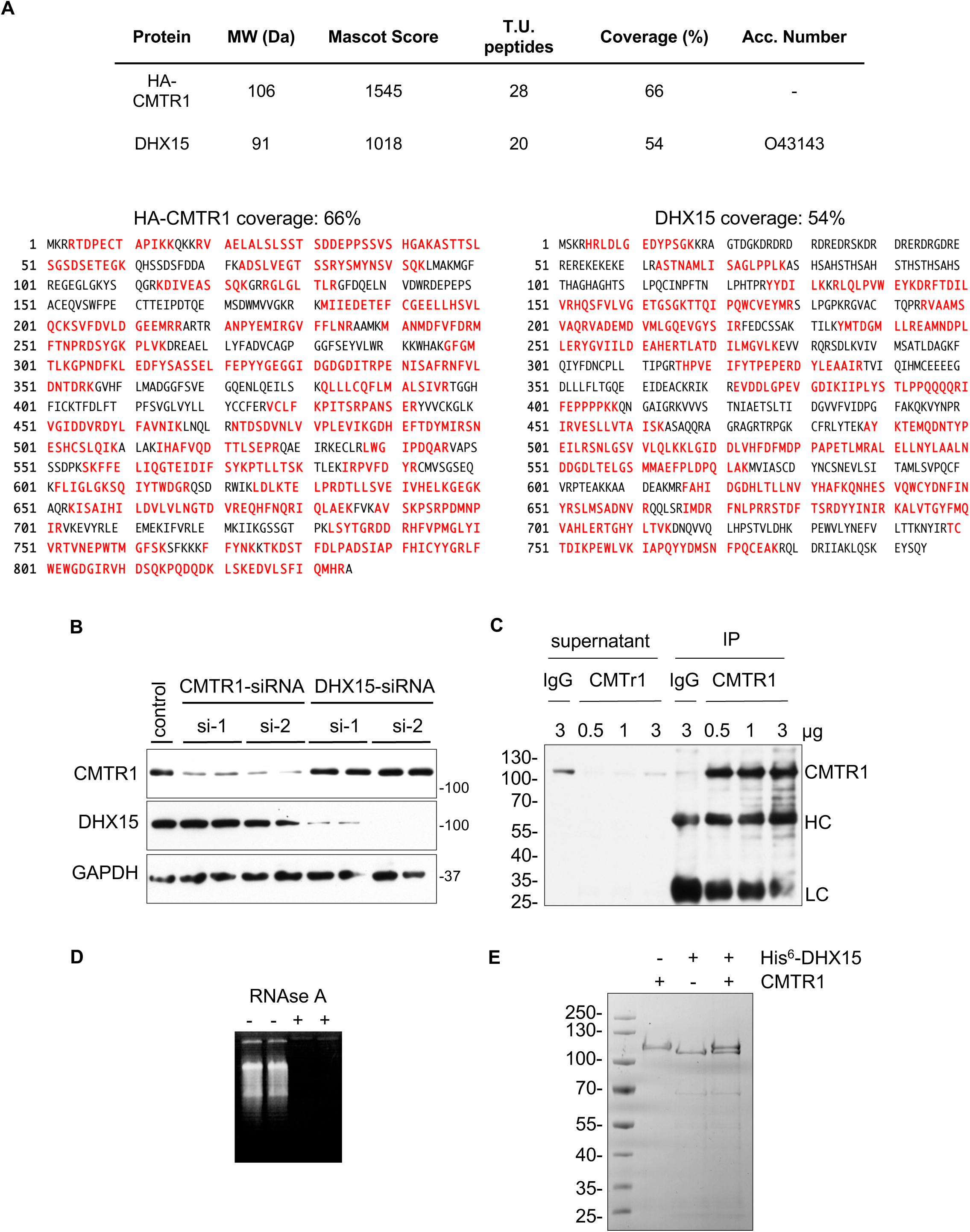
a) Mascot scores and protein coverage of HA-CMTR1 and DHX15. b) HeLa cells transfected with two independent CMTR1 or DHX15 siRNAs or non-targeting control. Cell extracts analysed by western blot. c) CMTR1 IP from HeLa cell extracts with indicated amount of antibody. Western blot analysis performed on supernatant or IPs. HC, heavy chain; LC, light chain. d) 2 μg HeLa RNA incubated with or without RNaseA and resolved by electrophoresis. e) 100ng recombinant His^6^-DHX15 and/or CMTR1 resolved by SDS-PAGE and Coomassie-stained.

**Figure EV3.**
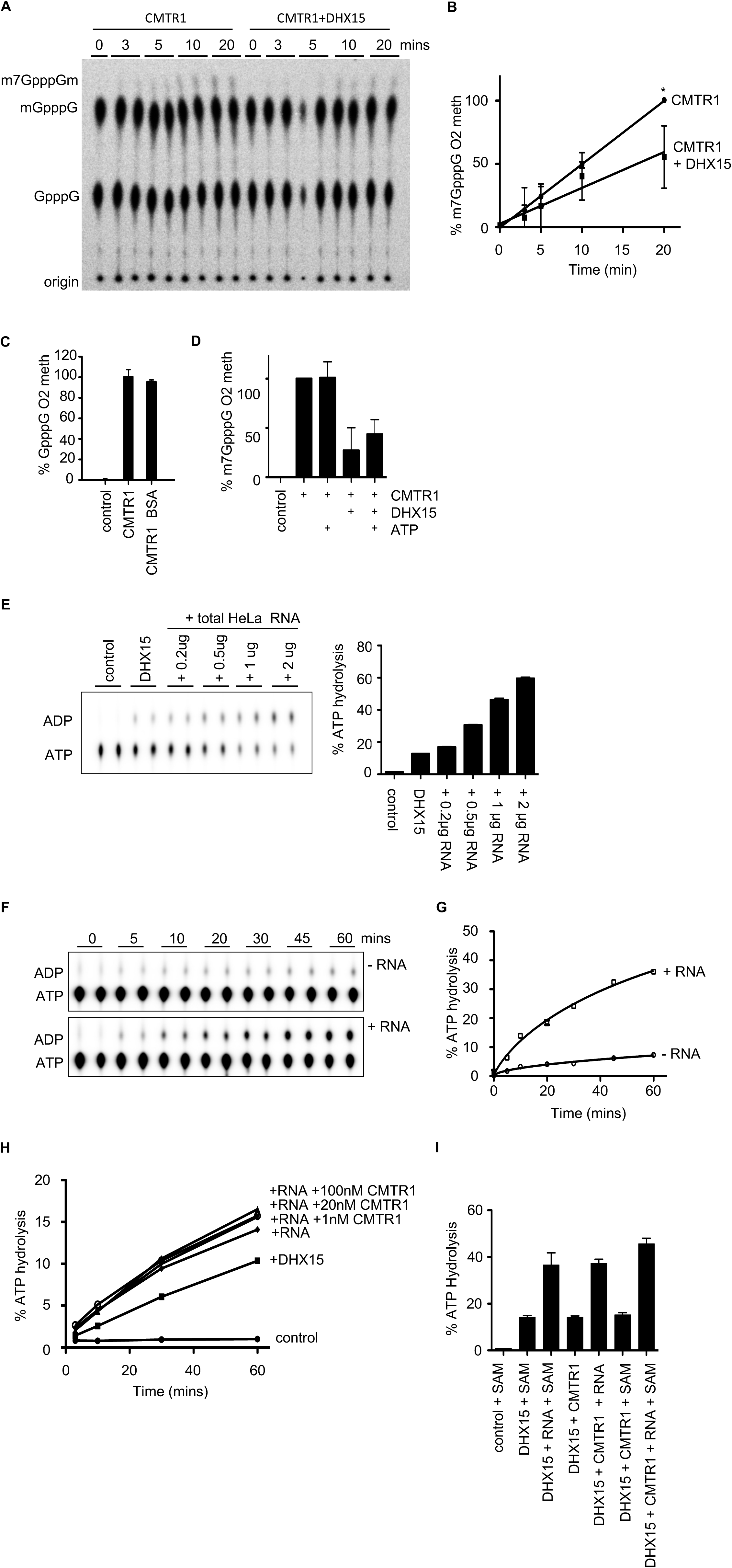
a) m7GpppG-capped RNA incubated with 3nM CMTR1, s-adenosyl methionine (SAM) and 3nM DHX15. Caps throughout figure ^32^P-labelled on α-phosphate. Generation of m7GpppGm (0-2 methylated) detected by thin layer chromatography (TLC) and autoradiography. b) Percentage conversion of m7GpppG to m7GpppGm plotted. Average and standard deviation for three independent experiments. Students T test performed. "*", P value<0.05. c) GpppG-capped RNA incubated with 3nM CMTR1, SAM and +/-3nM BSA. Generation of GpppGm (0-2 methylated) detected. d) m7GpppG-capped RNA incubated with 3nM CMTR1, SAM, +/-3nM DHX15, +/-300μM ATP. Generation of m7GpppGm (0-2 methylated) detected. e) 100nM DHX15 incubated with α^32^P-ATP and indicated HeLa RNA. Generation of ADP detected by TLC and autoradiography. f) As in (e) except reaction performed +/-1ug RNA for over a time course. g) Percentage ATP hydrolysis plotted with respect to time. h) 100nM DHX15 incubated with 1μg RNA and a titration of recombinant CMTR1. Percentage ATP hydrolysis plotted with respect to time. Control, no DHX15. i) As in (e) except 100nM DHX15 incubated with SAM, RNA and 100nM CMTR1 as indicated. Percentage ATP hydrolysis plotted.

**Figure EV4.**
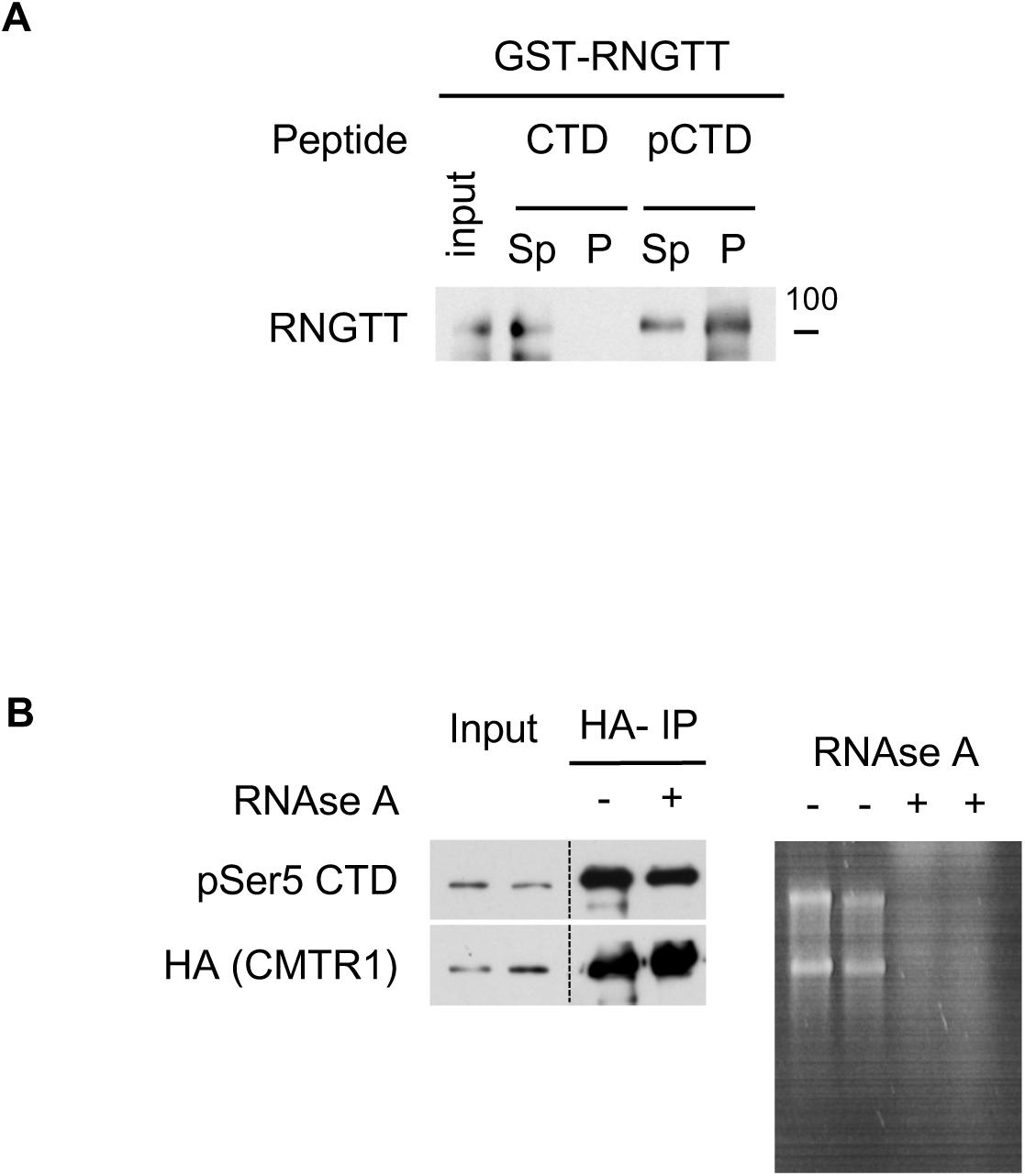
a) Recombinant GST-RNGTT incubated with biotinylated peptides consisting of 3 RNA pol II CTD heptad repeats, either unphosphorylated (CTD) or phosphorylated on S5 (pCTD). Peptides and co-purifying proteins enriched on streptavidin beads and western blot analysis performed. Aff, affinity purified fraction; FT, flow through. b) HA-CMTR1 IP from HeLa cell extracts, untreated (-) or incubated with RNAase A (+) before western blot analysis. In parallel, 2 μg HeLa RNA incubated with RNAase A and resolved on an agarose gel.

**Figure EV5.**
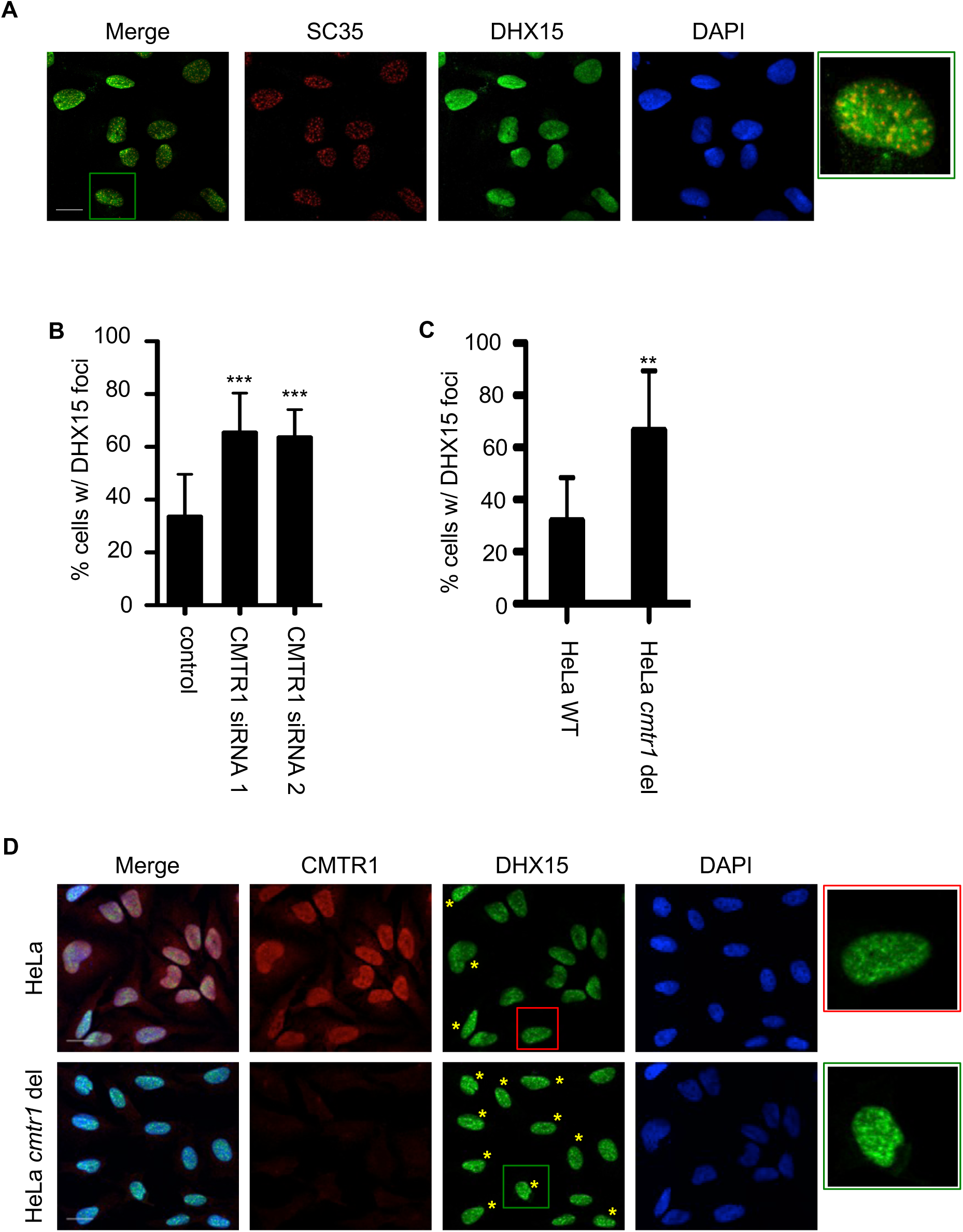
a) Immunofluorescence in HeLa cells. Individual and merged images show SC35 (red), DHX15 (green) and DAPI (blue). Quantification of number of cells with DHX15 foci in b) HeLa cells transfected with two independent CMTR1 siRNAs or control (a minimum of 175 cells were counted in 12 independent fields from two experiments per condition); and c) HeLa or HeLa *CMTR1* del cells (a minimum of 100 cells were counted in 9 independent fields from two experiments per condition). Average and standard deviation of cells with accumulated DHX15 foci within each field represented. Students T test performed. "**", P value<0.01; "***", P value<0.005. d) Representative field of endogenous CMTR1 and DHX15 immunofluorescence in HeLa and HeLa *CMTR1* del cells. Individual and merged images show CMTR1 (red), DHX15 (green) and DAPI (blue). Yellow asterisks indicate cells with DHX15 foci. 4.5x magnified areas are marked with coloured squares. Bar indicates 20μm.

